# Visualizing the Chronicle of Multiple Cell Fate using a Near-IR Dual-RNA/DNA-Targeting Probe

**DOI:** 10.1101/2025.05.29.655717

**Authors:** Linawati Sutrisno, Gary J. Richards, Jack D. Evans, Michio Matsumoto, Xianglan Li, Koichiro Uto, Shigehiro Yamaguchi, Jonathan P. Hill, Masayasu Taki, Katsuhiko Ariga

**Author notes:** Corresponding author. (L.S.); (J.P.H.); (M.T.); (K.A.).

## Abstract

Early detection and late-stage cell fate assessment are key factors to develop therapeutic strategies, although current methods cannot capture early responses or distinguish multiple injury states, especially in UV-vis-sensitive cells. Here, we introduce a method to simultaneously detect variations in RNA and DNA under near-infrared photoexcitation. Utilizing a pyrazinacene-based probe (TEG_8_-N14), we unexpectedly achieved discrimination of multiple cell states, including apoptosis, necrosis, necroptosis, and senescence, based on RNA–DNA changes. Specifically, TEG_8_-N14 selectively stains necrotic cells in live samples, while after fixation, it allows detection of ultra-early senescence in UV-vis-sensitive cells, providing approximately two-fold greater informational content than existing RNA or DNA fluorophores. These findings break current imaging barriers by enabling comprehensive visualization of single-cell fate histories without being affected by UV-vis or genetic manipulation.

## Introduction

Accurate monitoring of the fate of a single cell following the application of external stimuli is critical to the understanding of injury mechanisms and for the evaluation of therapeutic efficacy *(1)*. However, current approaches for identifying cell fate transitions in heterogeneous cell populations fail simultaneously to detect different possible biological processes often yielding inaccurate results, leading to an incomplete picture of cell fate post-treatment or erroneous conclusions in biological studies. For example, while caspase-3 is widely utilized as an apoptosis marker, it also participates in pathological processes, including oligodendrocyte injury in early multiple sclerosis lesions, confounding interpretation of imaging data *(2–3)*. Similarly, Acridine Orange (AO), one of the few dyes capable of detecting early-stage apoptosis and necrosis, exhibits high cytotoxicity and requires UV-vis excitation, rendering it incompatible with live or UV-vis-sensitive cells *(3–6)*. Nuclear morphology is regarded as a reliable, universal senescence marker, although it provides limited sensitivity due to minimal morphological variations during early stages of senescence *(7–10)*. An effective approach to assess single-cell fate transitions should meet the following criteria: (i) be operable in the NIR excitation range (≥640 nm) for broad applicability even in UV-vis-sensitive cells, (ii) enable early detection of cell injury states, (iii) be highly informative in distinguishing different forms of cell injury post-treatment, (iv) support multiplex imaging for analysis of complex biosystems, (v) exhibit superior sensitivity to cellular variations, and (vi) demonstrate exceptional photostability for prolonged imaging. Next-generation dyes should meet these criteria to enable accurate imaging results.

Here, we present a general strategy based on a class of *N*-heteroacene fluorophores, the pyrazinacenes, which have not so far been explored for use in bioimaging *(11–12)*. We have explored the importance of NIR excitation of the pyrazinacene chromophore for cellular imaging, and demonstrate the superiority of the probe over existing dyes (fig. S1 and 2, Fig. 1A). Specifically, pyrazinacene (TEG_8_-N14) enables simultaneous RNA-DNA imaging under NIR excitation. Computational modeling revealed distinct interaction modes that contribute to the differential spectral responses of the probe in RNA-versus DNA-rich environments. The superior DNA-RNA sensitivity and exceptional photostability of TEG_8_-N14 enable its application for precise detection of different states of cell injury (Fig. 1, B and C). To demonstrate this, we have used it to detect senescence in UV-vis-sensitive cells using autofluorescence-free channels (Fig. 1, D and E). Interestingly, our findings reveal that, compared to conventional nuclear morphology-based methods, RNA offers greater sensitivity for the prediction of cellular senescence. These findings establish a solid conceptual framework as a strategy to visualize chronicle of single-cell fate transitions, surpassing the limitations of any currently available imaging systems.

**Fig. 1.**
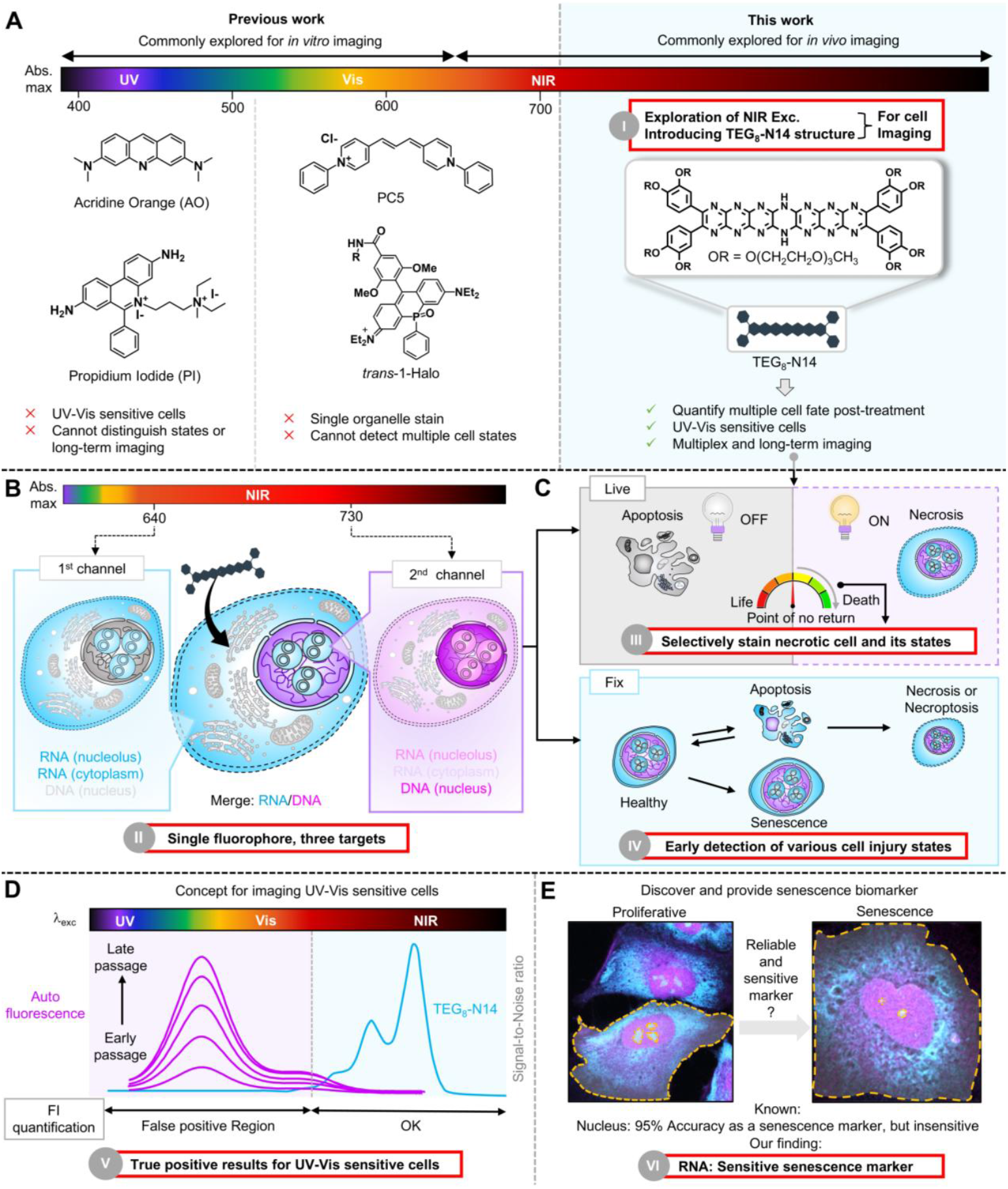
RNA-DNA and cell state discrimination using a single fluorophore under NIR excitation. (**A**) Comparison of TEG_8_-N14 and conventional cellular dyes for detection of various cell injury states. (**B**) Illustration of simultaneous RNA-DNA visualization using TEG_8_-N14 under dual NIR photoexcitation, enabling single-step staining for imaging complex biosystems and reducing the need for numerous control groups. (**C**) TEG_8_-N14 exhibits different imaging behavior in live and fixed cellular systems. In live cells, TEG_8_-N14 acts as a biomarker for irreversible necrosis, allowing for detailed, state-specific quantitative analysis of cell injury progression. In fixed cells, it facilitates precise discrimination among the states of apoptosis, necrosis, and necroptosis. (**D**) Our proposed concept to address the imaging barrier in UV-Vis sensitive cells. High autofluorescence and reactivity of endogenous fluorophores in ARPE-19 cells hinder the performance of existing UV-Vis dyes, highlighting the need for TEG_8_-N14 in FI quantification for this cell line. FI, Fluorescence Intensity. (**E**) Demonstration of RNA as sensitive cell stress biomarker in ARPE-19 cells using TEG_8_-N14.

### Introducing pyrazinacene to the bioimaging field

To address the requirements for cell fate imaging, we employed the tetradecaazaheptacene (N14) chromophore which exhibits strong absorption in the biological transparency window (650–900 nm) (*13*). For water solubility, triethylene glycol monomethyl ether appendages (*n* = 8) were introduced yielding hydrophilic TEG_8_-N14. Given the nitrogen-rich, planar structure of the N14 core, we hypothesized that TEG_8_-N14 might operate either as DNA groove binder or interact with RNA chains through hydrogen bonding (as donor or acceptor) or hydrophobic interactions *(13– 14)*. However, N14 chromophores are known to aggregate causing aggregation-induce fluorescence emission quenching (*12*). As expected, TEG_8_-N14 exhibits a peak absorbance in aqueous solution at around 664 nm and an extremely low quantum yield (below 0.01), caused by aggregation of the chromophores (fig. S3 and table S1). To modulate aggregation and enhance its emissive properties, TEG_8_-N14 was added to an aqueous cetyltrimethylammonium bromide (CTAB) solution, and used in subsequent comparison studies to investigate the effect of aggregation. In the presence of CTAB, pronounced enhancement of the 0-0 transition peak at approximately 730 nm was observed, accompanied by increases in molar absorption coefficient (εPBS buffer = 243,000 M−1 cm−1) and photoluminescence quantum yield (PLQY, ∼0.23), indicating that interactions with CTAB induce fluorescence switch-on of TEG_8_-N14 in aqueous solution due to changes in its state of aggregation. Spectral analysis reveals that these compounds are pH-sensitive (5.8–8.0) and remain stable at storage and application temperatures (figs. S4 and S5). Furthermore, we note that TEG_8_-N14 in the presence of CTAB is one of the brightest NIR dyes for cellular imaging among the 13 popular NIR dyes reported to date (*13–21*; table S1).

### RNA-DNA discrimination using a single fluorophore

Encouraged by the favorable optical properties of TEG_8_-N14, we evaluated its performance in cellular imaging. Cytotoxicity assays confirmed that TEG_8_-N14 is essentially non-toxic below 10 μM, and this condition was selected for further live cell imaging experiments (fig. S6). Unexpectedly, imaging of fixed HeLa cells treated with TEG_8_-N14 revealed distinct staining patterns when excited at 640 nm or 730 nm (fig. S7 and Fig. 2A). Based on subcellular localization, we envisioned that signals excited at 640 nm originate predominantly from RNA-rich regions, while signals at 730 nm mainly correspond to DNA-rich regions. This observation indicates that TEG_8_-N14 enables simultaneous staining of RNA and DNA in a single step. This dual labeling capability offers advantages for multiplexed imaging by reducing the number of control groups required. We further examined colocalization of TEG_8_-N14 with RNA and DNA by co-staining with a commercially available dye (Fig. 2B). Co-staining with Hoechst 33342 revealed the high DNA specificity of TEG_8_-N14 (R = 0.83 ± 0.06; *n* ≥ 21 cells), while RNA colocalization analysis yielded a Pearson correlation coefficient of 0.95 ± 0.01 (*n* ≥ 20 cells) with high overlap across all *z*-positions (fig. S8). The high specificity found here for RNA-DNA labeling is supported by low degrees of overlap with different organelles (fig. S9). This finding is further supported by fluorescence enhancement of TEG_8_-N14 obtained on the addition of commercially available dsDNA, ssDNA, and RNA, confirming their interaction with DNA and RNA within cellular systems (table S2, figs. S10 and S11).

**Fig. 2.**
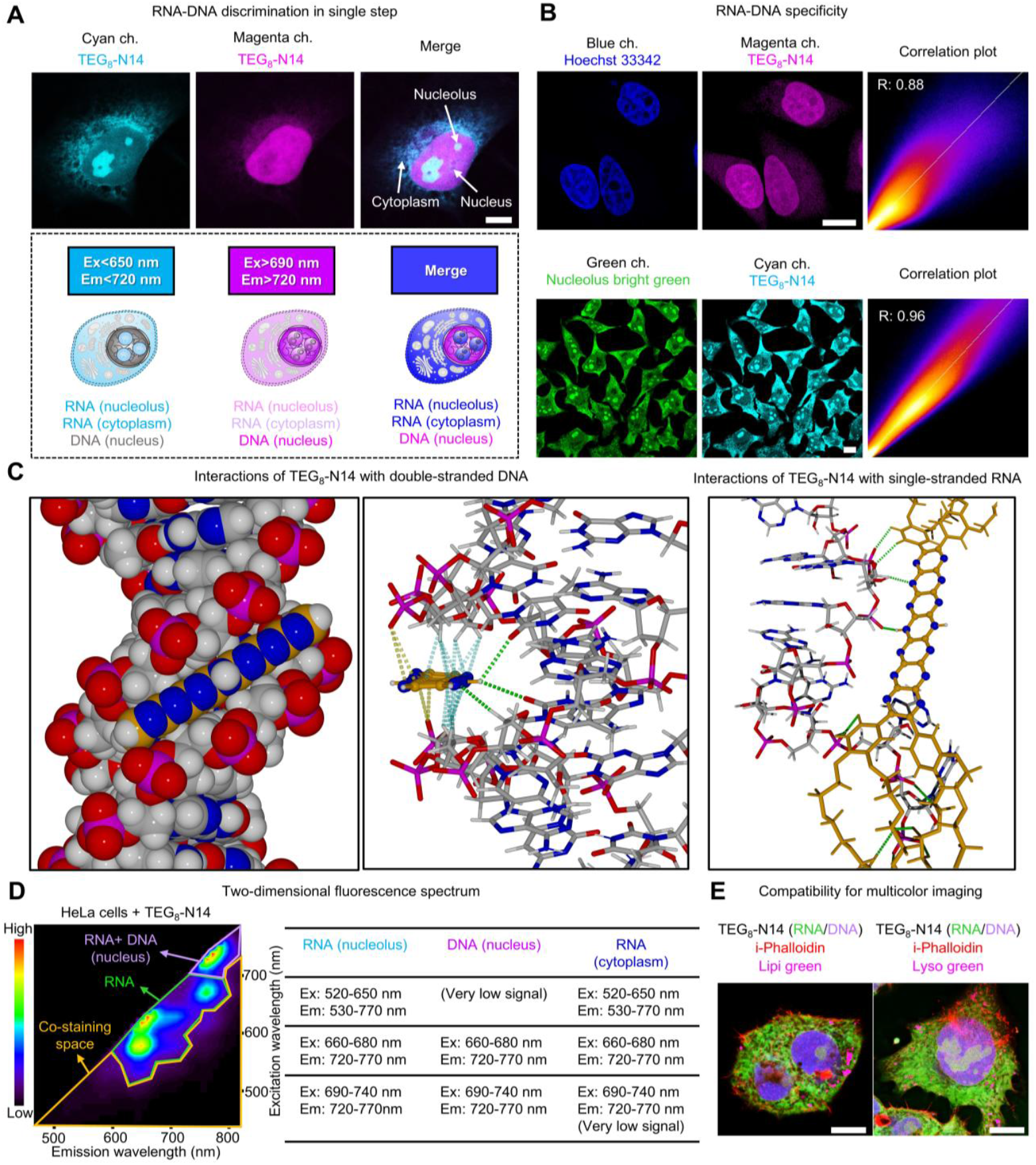
Performance of TEG_8_-N14 in fixed cell imaging. (**A**) Representative confocal images of HeLa cell stained with TEG_8_-N14 (upper) and proposed pseudo-color image of TEG_8_-N14-treated HeLa cells in different channels (lower). See movies S1–S4. Ex, excitation. Em, emission. Scale bar: 10 μm. (**B**) Representative co-staining images (left and middle) and their colocalization analysis (right). DNA colocalization is observed in blue (λ_ex_ = 405 nm and λ_em_ = 415–600 nm) and magenta (λ_ex_ = 730 nm and λ_em_ = 740–850 nm) channels. RNA colocalization is observed in green (λ_ex_ = 450 nm and λ_em_ = 500–550 nm) and cyan (λ_ex_ = 640 nm and λ_em_ = 650–720 nm) channels. Scale bar: 10 μm. Representative results of *n* = 3 over three independent experiments. Single focal plane. R is pearson correlation coefficient. See fig. S8 for three-dimensional colocalization. (**C**) Docking simulation to reveal interaction between TEG_8_-N14 with dsDNA (left and middle) and ssRNA (right). (**D**) Two-dimensional fluorescence imaging (left) and four-color imaging of fixed HepG2 cells stained with TEG_8_-N14 and several dyes (right). Red: i-Phalloidin (λ_ex_ = 405 nm, λ_em_ = 415–445 nm), magenta: Lipi green (λ_ex_ = 470 nm, λ_em_ = 500–600 nm) or Lyso green (λ_ex_ = 450 nm, λ_em_ = 500–600 nm), green: TEG_8_-N14 (λ_ex_ = 640 nm, λ_em_ = 650–720 nm), purple: TEG_8_-N14 (λ_ex_ = 730 nm, λ_em_ = 740–850 nm). The color scale bar indicates the fluorescence intensity ratio. Scale bars: 10 μm. Representative results of *n* = 3 over two independent imaging experiments. (**E**) Summary of excitation and emission wavelength for detection of TEG_8_-N14 for two targets (RNA (nucleolus and cytoplasm) and DNA (nucleus)). All the cells were preserved in a fixed state and captured using volumetric mode.

The molecular basis of the dual-binding behavior was investigated using docking simulations of TEG_8_-N14 with representative dsDNA, ssRNA, and dsRNA oligonucleotides. This revealed distinct interactions in a simulated implicit aqueous environment. The rigid planar structure of the N14 core preferentially inserts into the DNA minor groove (Fig. 2C; left and middle panels), where, one NH group of the N14 unit interacts with carbonyl atoms of two thymine residues by hydrogen bonding. There are also multiple C-H···π interactions *(22)* involving methylene units of ribose in the DNA minor groove. Close approach of phosphate oxygen atoms also suggests anion-π interactions involving the anionic phosphate backbone and electron deficient extremities of the N14 unit *(23)*. In contrast, interactions with RNA were more variable and dependent on its secondary structure. ssRNA undergoes hydrogen bonding with the N14 core and TEG side chains, while dsRNA binding is mediated largely by non-specific electrostatic interactions with the polyether appendages, with minimal π–π interaction (Fig. 2C, right; fig. S12). These simulations explain the observed spectral behaviors: binding to DNA significantly suppresses the geometry relaxation in the excited state, leading to enhanced 0-0 vibronic transition at ∼730 nm, while RNA interactions, which are more reliant on the aggregated state, promote increased 0–1 transition intensity (*24–25*; fig. S13).

Given its enhanced brightness, TEG_8_-N14 in the presence of CTAB was selected for multi-color imaging and photostability assay (fig. S14). Two-dimensional fluorescence TEG_8_-N14 demonstrating its ability to minimize crosstalk by opening up unpopular NIR excitation channel for multicolor imaging (Fig. 2, D and E, fig. S15, and movies S1–S4). We compared the photostability of TEG_8_-N14 with existing dyes based on half-lives of bleaching (t_1/2_ bleach) *(14)*. Photostability analysis shows that TEG_8_-N14 possesses excellent photostability comparable to SiR-DNA, arguably the most photostable dye reported to date, highlighting its potential for volumetric imaging (fig. S16 and table S3). RNA-DNA fluorescence signals were observed at the same localizations in different cell types and morphologies, demonstrating a broad applicability across various cell lines (fig. S17). To the best of our knowledge, this is the first dye that not only stains RNA and DNA simultaneously, but also possesses high photostability under NIR photoexcitation (≥640 nm) making it suitable for use studying diverse biological events.

### Broad applicability for measuring multiple single-cell injury states

Current microscopy techniques lack sensitivity to detect early apoptosis, rarely capture necrotic or necroptotic states, and typically require UV-vis excitation, limiting their applicability *(3)*. Given the role of RNA as an early injury marker, we demonstrate here the capability of TEG_8_-N14 to distinguish apoptosis, necrosis, and necroptosis. RNA sensitivity was first confirmed through strong fluorescence intensity attenuation upon RNase treatment (fig. S18), validating its responsiveness to RNA degradation. Spectral imaging of stained NIH-3T3 cells also confirms RNA and DNA detection in different channels, supporting RNA-DNA quantification for further study (fig. S19).

In order to evaluate the use of TEG_8_-N14 to distinguish different injury states, NIH-3T3 cells were treated with staurosporine (STS) to induce apoptosis, H_2_O_2_ to trigger necrosis, and TNF combined with a caspase inhibitor to provoke necroptosis *(3)*. In all cases, imaging after 6 hours revealed significant reduction in fluorescence intensities in both DNA and RNA channels (fig. S20). This result is consistent with the previous observation that nucleic acid degradation accompanies metabolic disruption during cell injury (*3, 26, 27*). Images were acquired at different times to identify distinct cell injury states using TEG_8_-N14 at the single cell level. Control cells exhibited flattened shapes with a cyan signal predominantly in the cytoplasm and a well-defined nucleus outlined fully by a magenta signal (fig. S21 and Fig. 3A). After 2 hours of STS treatment, most cells lose their flattened shape, develop thin extensions, undergo cellular fragmentation, and show less distinct, shrunken nuclei. At advanced states of apoptosis, which might correspond to onset of necrosis, the cyan signal in the cytoplasm is almost completely absent and nuclei are fragmented or show smooth and bright fluorescence intensity. In contrast to apoptosis, necrosis and necroptosis processes exhibit several distinct but mutually similar morphological features (Fig. 3B). At the early stages of these processes, there is a noticeable loss of cyan staining due to cellular shrinkage, although there is no cellular fragmentation. The initial signs include nuclear shrinkage and loss of distinct nuclear boundaries. At later stages of necrosis and necroptosis, nuclei shrink and often show increased intensity or a smooth appearance with minimal cytoplasmic cyan staining. The absence of both cytoplasmic and nuclear fragmentation—features characteristic of apoptosis—along with the presence of a smooth, round nucleus with low levels of cytoplasmic RNA, clearly differentiate necrosis and necroptosis from apoptosis. This highlights RNA as a highly sensitive indicator of early cell injury and supports the use of TEG_8_-N14 as a robust probe for discriminating multiple cell fates.

**Fig. 3.**
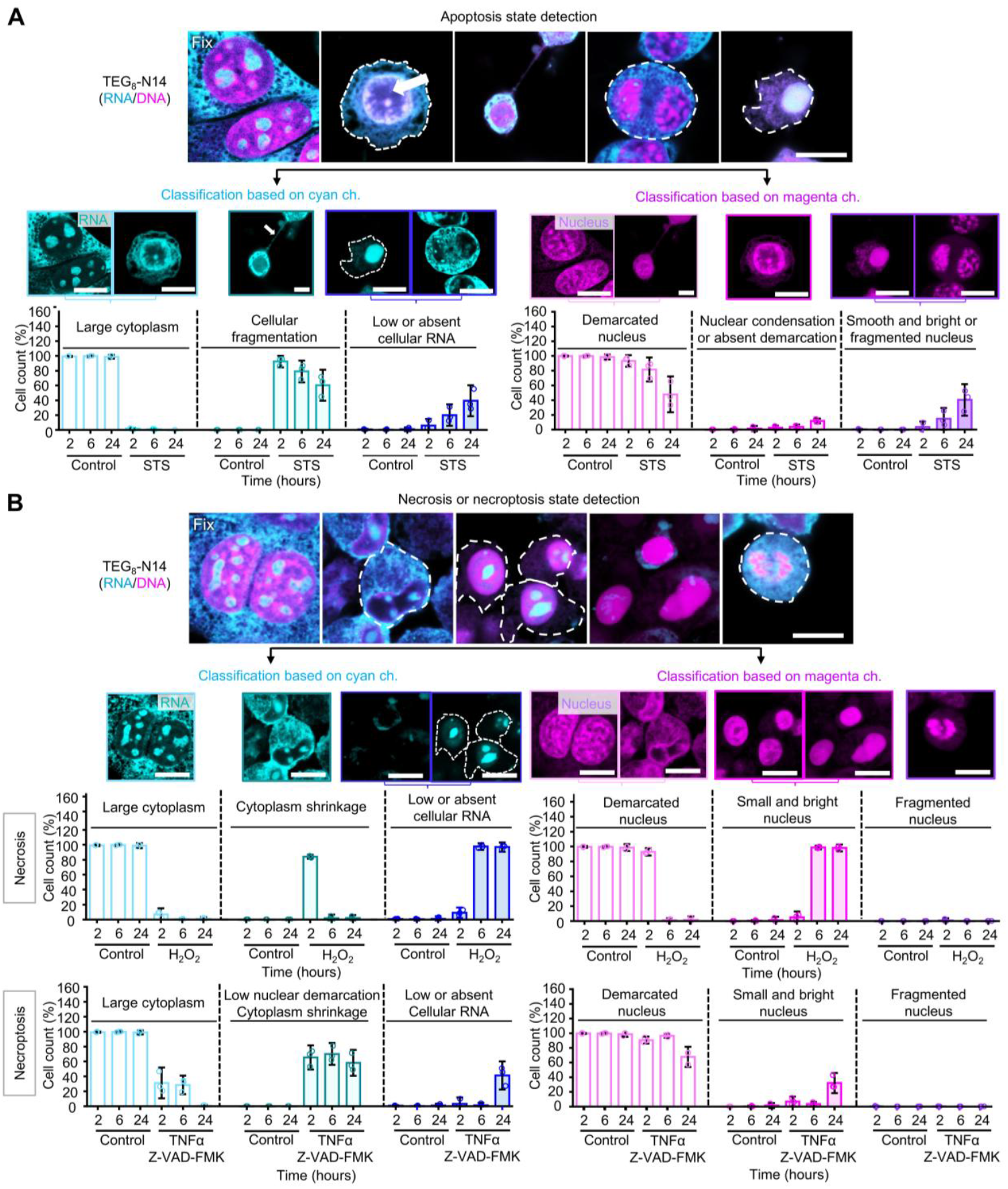
Single-cell analysis in different cell injury states by using TEG_8_-N14. (**A**) Representative confocal images of apoptotic states are shown (top), along with segmentation and analysis of apoptotic states based on cell morphology (bottom) in the cyan channel (λ_ex_ = 640 nm and λ_em_ = 650–720 nm) and magenta channel (λ_ex_ = 730 nm and λ_em_ = 740–850 nm). Scale bars: 10 μm. White arrow indicates nuclear condensation; white dashes indicate RNA in the cytoplasm. Data presented are mean ± S.D. (*n* > 250 cells per experiment). Dotted indicates three independent experiments. Error bars represent S.D. See fig. S21. STS, staurosporine. (**B**) Representative confocal images of necrosis and necroptosis states are shown (upper panels) and corresponding segmentation with analysis of necrosis and necroptosis states based on the morphology of cells (lower panels) in cyan channel (λ_ex_ = 640 nm and λ_em_ = 650–720 nm) and magenta channel (λ_ex_ = 730 nm and λ_em_ = 740–850 nm) are presented. Scale bars: 10 μm. White dashes indicate RNA in the cytoplasm. Data presented are mean ± S.D. (*n* > 250 cells per experiment). Dots indicate the three independent experiments. Error bars represent S.D. All the cells were preserved in a fixed state and captured using volumetric mode.

### Advantages of TEG_8_-N14 as a necrotic state indicator over conventional methods involving live cells

To evaluate the ability of TEG_8_-N14 to detect necrotic cells in live samples, we treated NIH-3T3 fibroblasts with STS or hydrogen peroxide to induce necrosis or mixed apoptosis and necrosis. (Fig. 4, A and B, fig. S22). Fluorescence emission from TEG_8_-N14 was observed only in cells with ruptured plasma membranes, thereby highlighting the potential of TEG_8_-N14 as a selective indicator of necrotic states. Further validation with an efflux pump inhibitor in live samples confirmed that TEG_8_-N14 cannot penetrate cells with intact plasma membranes (fig. S23), establishing that membrane rupture is required for probe entry. Traditionally, Propidium Iodide (PI) and AO have been gold standards for necrosis detection. However, both suffer from limitations: PI intercalates into DNA exhibiting long-term toxicity, while AO requires UV-vis excitation inducing phototoxicity (*28–29*), which restrict its use in live cells. In contrast to PI, which cannot be used to distinguish necrotic subtypes, TEG_8_-N14 allows identification of necrotic states based on RNA sensitivity (Fig. 4C). Moreover, in comparison with AO, TEG_8_-N14 exhibits ∼100-fold lower cytotoxicity, allowing continuous imaging up to 24 hours coupled with ability to detect necrotic cells (Fig. 4D and fig. S24). Overall, these results demonstrate that TEG_8_-N14 outperforms conventional necrosis markers by providing selective detection of necrotic states, minimizing cytotoxicity, and enabling long-term and reliable tracking of cell death in live-cell imaging experiments.

**Fig. 4.**
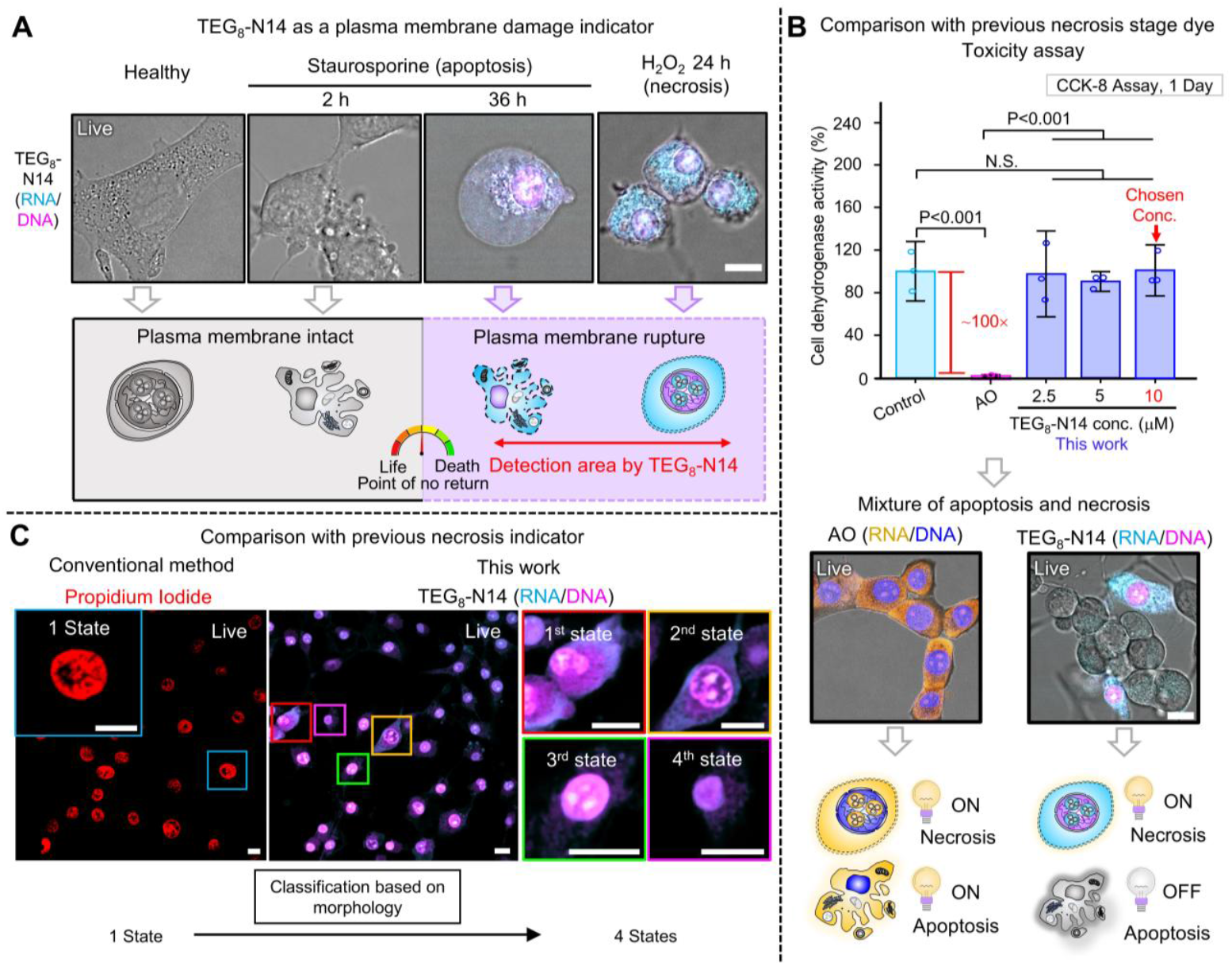
Comparison between TEG_8_-N14 with conventional necrosis dyes in live samples. (**A**) Representative confocal images of TEG_8_-N14 as necrosis indicator in NIH-3T3 cells. Images were obtained in cyan (λ_ex_ = 640 nm and λ_em_ = 650–720 nm) and magenta (λ_ex_ = 730 nm and λ_em_ = 740– 850 nm) channels. Scale bar: 10 μm. (**B**) Comparison of existing commercial RNA-DNA dye (Acridine Orange, AO) and TEG_8_-N14 for the detection of necrosis state in terms of cytotoxicity (upper) and necrosis detection (lower). Data presented as mean ± S.D. (*n* = 3). Error bars represent S.D. Significance levels are determined using an unpaired two-tailed Student’s t-test. Images were acquired in blue (λ_ex_ = 457 nm and λ_em_ = 467–550 nm), yellow (λ_ex_ = 457 nm and λ_em_ = 600–750 nm), cyan (λ_ex_ = 640 nm and λ_em_ = 650–720 nm) and magenta (λ_ex_ = 730 nm and λ_em_ = 740–850 nm) channels. Scale bar: 10 μm. (**C**) Comparison of existing necrosis indicator dye (Propidium Iodide, PI) and TEG_8_-N14 for the detection of necrosis state. Left image: red color indicates PI (plasma membrane damage indicator). Right images: cyan color indicates RNA stained with TEG_8_-N14 and magenta color indicates nucleus stained with TEG_8_-N14. Scale bars: 10 μm. All cells were directly stained without any fixative and imaged using volumetric mode.

### Discover sensitive biomarker in senescence using accurate detection method

Next, we explored the possibility of extending our classification strategy to detect cell senescence *(7, 30–34)*, a biological phenomenon whose detection is challenging. For this purpose, ARPE-19 cells were selected as a model because of their high level of endogenous fluorophores, which obstruct high reproducibility imaging *(5, 35–37)*. ARPE-19 cells were treated with doxorubicin (Dox) inducing ≥95% senescence *(38–40),* as verified by Trypan Blue assay (Fig. 5A). Prior to cell imaging, ARPE-19 cells were imaged using different excitation wavelengths to identify the optimum channel for observation. As expected, for excitation below 640 nm, ARPE-19 cells exhibited autofluorescence which increased at higher passage numbers of the cells (Fig. 5B). In contrast, when cells were imaged under excitation above 640 nm even at the maximum laser power of the microscope, almost no autofluorescence was observed confirming that NIR excitation channels are suitable for the accurate quantification of fluorescence intensity (FI) of this cell line.

**Fig. 5.**
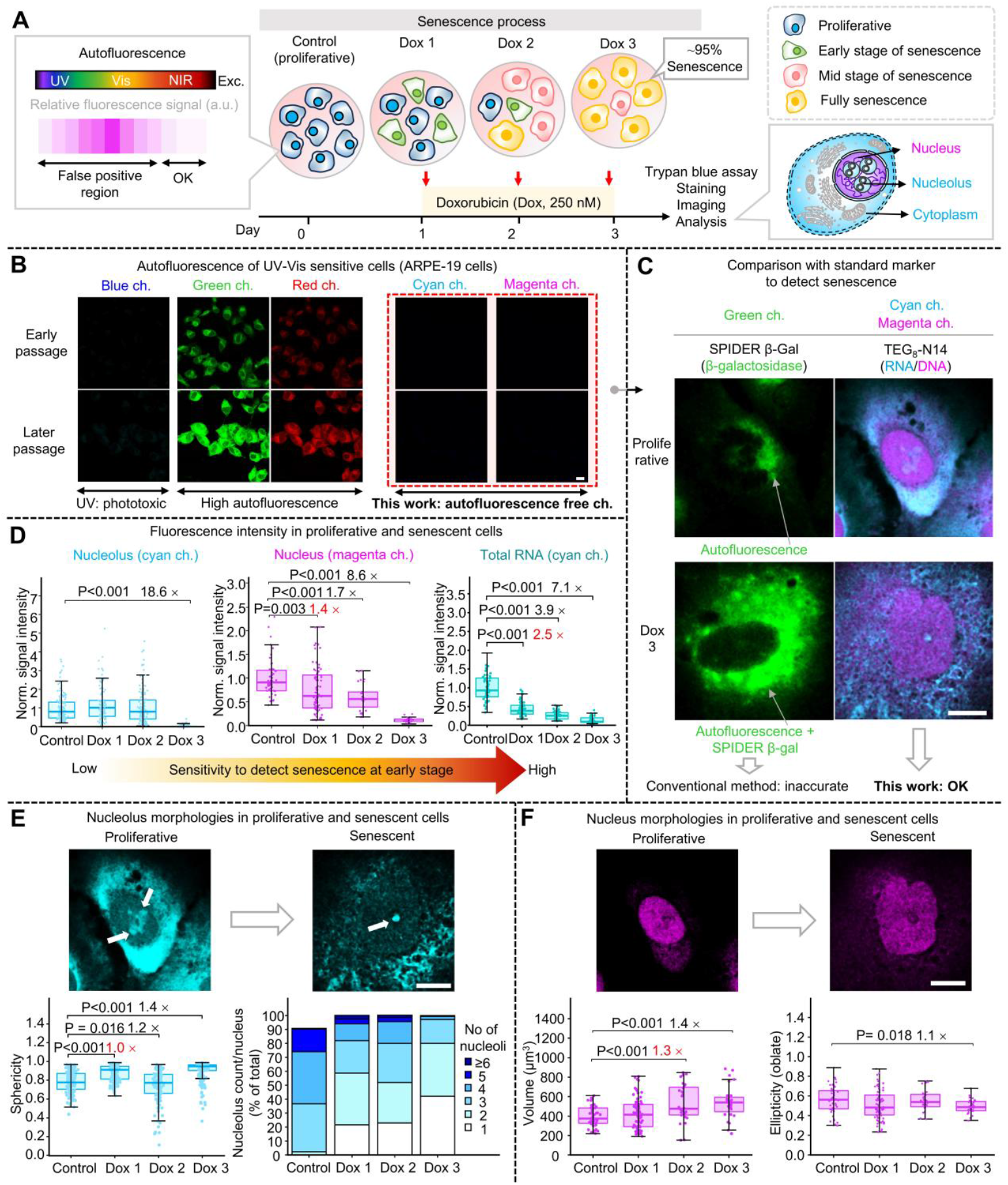
Reliable measurements in senescence-induced UV-vis sensitive cells by TEG_8_-N14. (**A**) Schematic illustrations of chemically-induced senescent cells amongst ARPE-19 cells and its imaging process. Exc.: excitation. (**B**) Representative confocal images of maximum intensity projection of endogenous fluorophores in ARPE-19 cells under different excitation conditions with identical microscope setup. Blue channel: λ_ex_ = 405 nm, λ_em_ = 415–460 nm, green channel: λ_ex_ = 488 nm, λ_em_ = 500–550 nm, red channel: λ_ex_ = 561 nm, λ_em_ = 571–635 nm, cyan channel: λ_ex_ = 640 nm, λ_em_ = 650–720 nm, magenta channel: λ_ex_ = 730 nm, λ_em_ = 740–850 nm. Scale bar: 10 μm. Representative results for *n* = 3 over two independent experiments. (**C**) Representative confocal images of proliferative and senescent ARPE-19 cells stained with SPIDER β-gal and TEG_8_-N14. Merged images were obtained in green (λ_ex_ = 488 nm, λ_em_ = 500–550 nm), cyan (λ_ex_ = 640 nm and λ_em_ = 650–720 nm) and magenta (λ_ex_ = 730 nm and λ_em_ = 740–850 nm) channels. See fig. S26 for low magnification images. (**D**) Fluorescence intensity changes in nucleolus (left), nucleus (middle), and total RNA (right) measured after treatment with Dox to induce senescence from day 1 to day 3. Data are presented as mean ± S.D. and plotted from three biological replicates, *n* > 30 cells. Error bars represent S.D. Statistical significance was assessed using an unpaired two-tailed t-test, n.s.: non-significant. Dox, doxorubicin. Three-dimensional morphological images (upper) and its analysis (lower) of the nucleolus (**E**) and nucleus (**F**). Data are presented as mean ± S.D. and plotted from three biological replicates, *n* > 30 cells. White arrows indicate nucleolus. Error bars represent S.D. Scale bar: 10 μm. Statistical significance was assessed using an unpaired two-tailed t-test, n.s.: non-significant. Dox, doxorubicin. All cells were treated with a fixative and imaged using volumetric mode.

Senescence in ARPE-19 cells is difficult to detect using conventional RNA/DNA dyes due to UV-vis excitation requirements, endpoint limitations, and signal crosstalk. Although nuclear morphology is ∼95% accurate, its low early-stage sensitivity highlights the need for better biomarkers *(3, 7, 41-42)*. With the autofluorescence-free properties of TEG_8_-N14 in mind, we simultaneously visualized RNA in the nucleolus, RNA in the cytoplasm, and the nuclear DNA. First, we compared our system with the conventional senescence marker SPIDER β-gal (Fig. 5C and fig. S25). As expected, SPIDER β-gal imaging was subject to significant interference involving crosstalk between the dye and endogenous fluorophores in ARPE-19 cells. In contrast, TEG_8_-N14 gave more accurate results for the easy detection of senescence due to the autofluorescence-free channel (Fig. 5, B and C). To the best of our knowledge, these results represent the first RNA-DNA visualization of chemically-induced senescence in ARPE-19 cells using dual autofluorescence-free channels.

Based on these results, we conducted an in-depth analysis using our system (fig. S26 and Fig. 5D). FI quantification showed a ∼2.5-fold decrease in total RNA, surpassing changes in the nucleolus and nucleus at the earliest senescence state, indicating total RNA as a promising and sensitive senescence marker. Next, we performed quantitative analyses of morphological changes in the nucleus and nucleolus using volumetric segmentation analysis (Fig. 5, E and F, figs. S27 and S28). We observed that the sphericity of the nucleolus increased, while the nucleolus counts per cell decreased, aligned with previous findings *(39)*. Additionally, the nucleus became larger and more flattened after the cells entered senescence, consistent with previous results *(7)*. These findings imply that FI quantification of total RNA using TEG_8_-N14 is the most sensitive method for detecting senescence in ARPE-19 cells, surpassing changes in the nucleolus and nucleus at the earliest states of senescence. TEG_8_-N14 outperforms individual RNA or DNA dyes since it provides ∼2-fold higher information level, has superior RNA sensitivity, eliminates the requirement for UV-vis excitation, and can be used even in ARPE-19 cells. Our method also enables the distinction of senescence by combining a widely regarded reliable marker (nuclear morphology) with a sensitive marker (RNA) in a single step, while providing accurate FI quantification.

## Discussion

Precise determination of cell fate transitions in heterogeneous populations requires probes capable of sensitively and simultaneously monitoring multiple molecular states at the single-cell level *(3, 44–47)*. While both RNA and DNA play central roles in defining cellular states, conventional nucleic acid dyes are typically limited to a single target, rely on UV–vis excitation, and often suffer from phototoxicity and autofluorescence, particularly in sensitive or highly autofluorescent cell lines (*4, 13*). Here we demonstrate the usefulness of pyrazinacene dye, TEG_8_-N14, effectively overcoming the imaging barriers of conventional fluorophores. Our results highlight six key advantages of TEG_8_-N14 over conventional RNA-DNA stains. These are: (**1**) its unique dual-staining capability enables the simultaneous visualization of RNA and DNA (Fig. 1 and Fig. 2B). This feature enhances usability by reducing the need for multiple control groups and provides a high informational level for the monitoring of biological events. (**2**) By eliminating the need for UV-vis excitation, TEG_8_-N14 avoids phototoxicity, autofluorescence, and spectral overlap, allowing its application in UV-vis-sensitive cells (Fig. 2D and Fig. 5B). (**3**) It exhibits high photostability (fig. S16) for quantitative fluorescence imaging without photobleaching artifacts. (**4**) Its RNA sensitivity enables the detection of a wide range of cell injury states, from the earliest to the later states, including apoptosis, necrosis, and necroptosis (Fig. 3). (**5**) In live cell imaging, TEG_8_-N14 selectively accumulates in necrotic cells, providing a reliable platform for real-time necrosis tracking (Fig. 4). In contrast to PI and AO, which suffer from high cytotoxicity and low specificity, TEG_8_-N14 allows for long-term imaging up to 24 h. (**6**) In ARPE-19 cells, TEG_8_-N14 offers autofluorescence-free quantification of RNA and nuclear morphology, outperforming standard senescence markers for sensitivity and resolution (Fig. 5).

Overall, these advantages position TEG_8_-N14 as a next-generation multifunctional probe. Beyond its technical performance, our findings demonstrate that RNA is a highly sensitive and functional biomarker for cellular stress. Owing to its rapid turnover, high susceptibility to degradation, and greater structural flexibility *(3, 29)*, RNA reflects metabolic perturbations earlier than DNA. Coupling with morphological analysis of the nucleus through different NIR channels enables multimodal readouts of complex cell states.

From a fundamental viewpoint, our study marks an important milestone in the development of multifunctional fluorophores capable of analyzing distinct biological processes with molecular precision. From a practical application perspective, TEG_8_-N14 simplifies imaging workflows by reducing the need for multiple control groups, minimizing cytotoxicity, and avoiding UV-vis excitation. These features make it particularly suited for applications in live cell assays and high throughput drug screening. Looking ahead, advancing the understanding of pyrazinacene structure-function relationships and how they relate to biological phenomena is expected to drive substantial progress in both scientific research, industrial, and medical applications. Broadening the scope of TEG_8_-N14 and its derivatives to *in vivo* systems could open new frontiers in single cell diagnostics, spatiotemporal mapping of cell fate transitions, and preclinical evaluation of therapeutic efficacy.

## ACKNOWLEDGEMENTS

We thank Z. Luo (Chongqing University), S. Kewei (NIMS), O. Suguru (Leica, Co., Ltd), Y. Yamashita (NIMS) for valuable discussions during the preliminary stages of this project. We are also grateful to M. Ebara (Univ. of Tsukuba), N. Shirahata (NIMS), T. Nakanishi (NIMS), Z. Guo (NIMS) and ICYS administrative office (NIMS) for providing facilities and assistance during experiments. J.D.E (Univ. of Adelaide) is the recipient of an Australian Research Council Discovery Early Career Award (Project Number: DE220100163) funded by the Australian Government. Phoenix HPC service at the University of Adelaide provided high-performance computing resources.

## Author contributions

L.S. conceived the idea of this project. L.S., M.T., J.P.H., and K.A. planned and supervised the project. L.S., M.M., and M.T. designed the experiments. L.S. performed the characterization and conducted all cell biology experiments. G.J.R. conceived the design of the NIR chromophore, performed its synthesis, and characterization with assistance on purification from J.P.H. L.S. performed the titration and quantum yield measurements. L.X.L. and M.T. contributed to cell imaging. L.S. and M.T. analyzed the imaging data. J.D.E. performed computational studies. L.S., G.J.R., M.T., and J.P.H. wrote the manuscript with the input from M.M., K.U., S.Y., and K.A.

## Data and materials availability

All data are available in the manuscript or supplementary materials.

## REFERENCESAND NOTES

1. M.C. Huppertz et al., Science 383, 890–897 (2024).

2. M. Shibata et al., J. Clin. Invest. 106, 643–653 (2000).

3. J. R. Plemel et al., J. Cell. Biol. 216, 1163–1181 (2017).

4. P.P. Laissue et al., Nat. Method 14, 657–661 (2017).

5. A. Lakkaraju, S. C. Finnemann, E. Rodriguez-Boulan, Proc. Natl Acad. Sci. USA 104, 11026– 11031 (2007).

6. A. Espagne et al., Nat. Biotechnol., 42, 872–876 (2024).

7. I. Heckenbach et al., Nat. Aging 2, 742–755 (2022).

8. S. Lee, C. A. Schmitt, Nat. Cell Biol. 21, 94–101 (2019).

9. C.H. Hsu, S.J. Altschuler, L.F. Wu, Cell, 178, 361–373 (2019).

10. V. Gorgoulis, et al., Cell 179, 813–827 (2019).

11. G. J. Richards, J. P. Hill, Acc. Chem. Res. 54, 3228–3240 (2021).

12. G. J. Richards et al., J. Am. Chem. Soc. 141, 19570–19574 (2019).

13. K. Uno, N. Sugimoto, Y. Sato, Nat. Commun. 12, 2650 (2021).

14. N. C. Shaner et al., Nat. Methods 5, 545–551 (2008).

15. S. Usama et al., J. Am. Chem. Soc. 145, 14647™14659 (2023).

16. G. Lukinavicius et al., J. Am. Chem. Soc. 138, 9365™9368 (2016).

17. Y. Koide et al., J. Am. Chem. Soc. 133, 5680–5682 (2011).

18. J. Bucevičius et al., Nat. Commun. 14, 1306 (2023).

19. L. G. Wang et al., Nat. Chem. 15, 729–739 (2023).

20. M. Grzybowski et al., Angew. Chem. Int. Ed. 57, 10137–10141 (2018).

21. Q. Wu et al., Angew. Chem. Int. Ed. Engl. 63, e202400711 (2024).

22. M.T. Blázguez-Sánchez et al., Chem. Eur. J. 20, 17640–17652 (2014)

23. B. L. Schottel, F. T. Chifotides, K. R. Dunbar, Chem. Soc. Rev. 37, 68–83 (2008).

24. P. Mondal, et al., Adv. Mater. Interfaces 9, 2200209 (2022).

25. V. Conti Nibali, et al., J. Phys. Chem. Lett. 11, 4809–4816 (2020).

26. D. Bertheloot, E. Latz, B. S. Franklin, Cell. Mol. Immunol. 18, 1106–1121 (2021).

27. S. Nagata, H. Nagase, K. Kawane, N. Mukae, H. Fukuyama, Cell Death Differ. 10, 108–116 (2003).

28. K. E. Zulauf, J. E. Kirby, Proc. Natl Acad. Sci. USA 117, 29839–29850 (2020).

29. I.U. Cevik, T. Dalkara, Cell Death Differ. 10, 928–929 (2003).

30. J. Nehme et al., Nat. Aging 4, 771–782 (2024).

31. E. Hara, Nat. Cell Biol. 26, 176–176 (2024).

32. I. Duran et al., Nat. Aging 4, 1167–1170 (2024).

33. I. Duran et al., Nat. Commun. 15, 1041 (2024).

34. F. Debacq-Chainiaux et al., Nat. Prot. 4, 1798–806 (2009).

35. J. Zhou, Y. P. Jang, S. R. Kim, J. R. Sparrow, Proc. Natl Acad. Sci. USA 103, 16182–16187 (2006).

36. T. Taubitz et al., eBioMedicine, 48, 592–604 (2019).

37. M. Ogrodnik et al., Cell, 187, 16, 4150 – 4175 (2024).

38. J.B. Chae et al., Geroscience 43, 2809–2833 (2021).

39. M.C. Lanz et al., Mol. Cell, 82, 3255–3269 (2022).

40. S. Victorelli et al., Nature, 622, 627–636 (2023).

41. M. Collado, M.A. Blasco, M. Serrano, Cell, 130, 223–233 (2007).

42. V. Gorgoullis et al., Cell, 179, 813–827 (2019).

43. M. Jo et al., Redox Biol., 75, 103279 (2024).

44. X. Wen et al., Nature 628, 648–656 (2024).

45. M.P. Baar et al., Cell 169, 132–147 (2017).

46. S. He, N.E. Sharpless, Cell, 169, 1000–1011 (2017).

47. S. Lee, C.A. Schmitt, Nat. Cell Biol., 21, 94–101 (2019).

